# Characterizing transcriptional heterogeneity through pathway and gene set overdispersion analysis

**DOI:** 10.1101/026948

**Authors:** Jean Fan, Neeraj Salathia, Rui Liu, Gwen Kaeser, Yun Yung, Joseph Herman, Fiona Kaper, Jian-Bing Fan, Kun Zhang, Jerold Chun, Peter V. Kharchenko

## Abstract

Single-cell transcriptome measurements are being applied at rapidly increasing scales to study cellular repertoires underpinning functions of complex tissues and organs, including mammalian brains. The transcriptional state of each cell, however, reflects a variety of biological factors, including persistent cell-type specific regulatory configurations, transient processes such as cell cycle, local metabolic demands, and extracellular signals. Depending on the biological setting, all such aspects of transcriptional heterogeneity can be of potential interest, but detecting complex heterogeneity structure from inherently uncertain single-cell data presents analytical challenges. We developed PAGODA to resolve multiple, potentially overlapping aspects of transcriptional heterogeneity by identifying known pathways or novel gene sets that show significant excess of coordinated variability among the measured cells. We demonstrate that PAGODA effectively recovers the subpopulations and their corresponding functional characteristics in a variety of single-cell samples, and use it to characterize transcriptional diversity of neuronal progenitors in the developing mouse cortex.

## Introduction

Single-cell transcriptome measurements provide an unbiased approach for studying the complex cellular compositions inherent to multicellular organisms. Increasingly sensitive single-cell RNA-sequencing (scRNA-seq) protocols developed over the last few years can detect up to 48% of mRNA molecules contained in an individual cell, and examine thousands of cells in parallel^1,2^. Analyses of cell-to-cell transcriptome differences in such data have been used to catalogue cell types comprising natural tissues^3–6^, reconstruct molecular timecourses of developmental progressions^7–12^, and identify characteristic subpopulations in human cancers^13,14^. Nevertheless, the analysis of such single-cell RNA-seq datasets remains challenging, as these measurements expose numerous differences between cells, only some of which may be relevant for the system-level functions.

A large proportion of the observed cell-to-cell variation can be attributed to technical noise stemming from processing of low amounts of starting RNA in each cell^15^. Similarly, some of the variation reflects low-level stochastic processes such as transcriptional bursting^16^. These effects pose difficulties for principal component analysis (PCA) and other dimensionality reduction approaches. In particular, non-Gaussian noise and the presence of outliers decrease the sensitivity of PCA, and increase the difficulty of assessing the statistical significance of the detected features. Even when cell-to-cell variation reflect prominent biological processes taking place within the measured population, these processes may not be of primary interest. For example, differences in the metabolic state or cell cycle stage may be common to multiple cell types, and can mask more subtle cell-to-cell variability associated with cell differentiation or other processes being studied^12^. Such cross-cutting transcriptional features represent alternative ways to partition the cell population, and pose a challenge for the commonly-used clustering approaches that aim to reconstruct a single subpopulation structure^3,5,6,12^.

Here we propose an alternative approach for analysis of transcriptional heterogeneity, based on statistical evaluation of coordinated expression variability of previously-annotated or automatically-detected gene sets. Gene set testing with methods such as GSEA^17^ has been extensively utilized in the context of differential expression analysis to increase statistical power and uncover likely functional interpretations. Similar rationale can be applied in the context of heterogeneity analysis. For example, while cell-to-cell variability in expression of a single neuronal differentiation marker such as *Neurod1* may be too noisy and inconclusive, coordinated upregulation of a dozen genes associated with neuronal differentiation in the same subset of cells would provide a prominent signature distinguishing a subpopulation of differentiating neurons. The Pathway And Geneset OverDispersion Analysis (PAGODA) described here relies on cell-specific error models to estimate residual gene expression variance, and identifies pathways and gene sets that exhibit statistically significant excess of coordinated variability (overdispersion). By testing many different pathways and gene sets, PAGODA is able to capture multiple, potentially intersecting, aspects of heterogeneity, and provides a non-redundant overview of the heterogeneity structure within the measured cell population (Figure 1).

**Figure 1.**
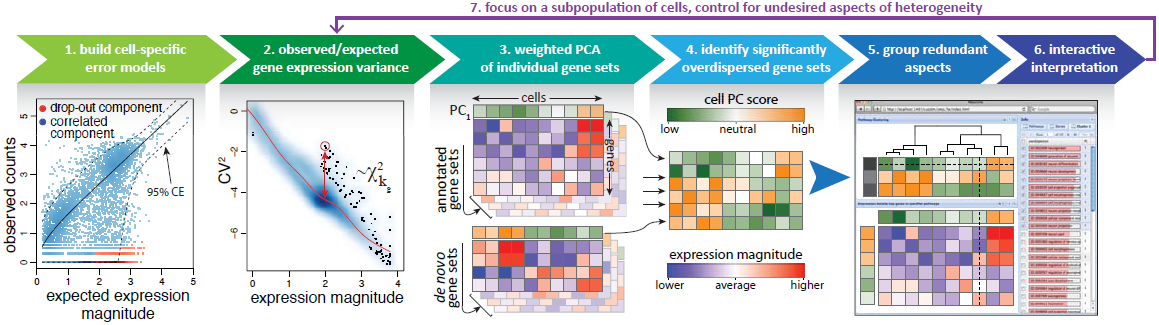
Pathway and gene set overdispersion analysis (PAGODA). The developed approach summarizes significant aspects of transcriptional heterogeneity through the following key steps: **1.** Parameters of probabilistic error models are determined for each cell to quantify the dependency of amplification noise and drop-out probabilities on the expression magnitude. The plot shows the model fit for a particular cell, separating drop-out and amplified (correlated) components, and the 95% confidence envelope of the amplified component based on the determined negative binomial dispersion dependencies; **2.** The magnitude of expression variance for each gene is rescaled relative to the transcriptome-wide expectation model (red curve), taking into account the uncertainty in the variance estimates of each gene by determining effective degrees of freedom (*kg*) for the χ^2^ distribution; **3.** Weighted PCA analysis is performed independently on functionally-annotated gene sets, as well as *de novo* gene sets determined based on correlated expression in the current dataset. The annotated gene sets increase statistical power to detect relevant aspects of heterogeneity and provide functional interpretation to the observed aspects of heterogeneity, whereas *de novo* gene sets can detect aspects of heterogeneity not captured by the annotated gene sets; **4.** If the amount of variance explained by a principal component of a given gene set is significantly higher than expected (see Methods), the gene set is called overdispersed, and the cell scores defined by that principal component (coded in orange-green gradient) are included as one of the significant aspects of heterogeneity in the dataset; **5** Redundant aspects that are driven by the same sets of genes or show very similar patterns of cell separation are grouped to provide succinct overview of heterogeneity; **6.** PAGODA implements a web browser-based interface to navigate the identified aspects of heterogeneity, associated gene sets and gene expression patterns. **7.** Depending on the biological question, some of the detected aspects of heterogeneity may be deemed artifactual or irrelevant, and can be actively controlled for in a subsequent iteration. Subsequent iteration can also be used to focus on a subpopulation defined by one or more of the detected aspects.

Single-cell analyses have been particularly useful for elucidating the complexity of neural tissues. While the extensive heterogeneity of neuronal morphologies has been recognized since the studies of Cajal^18^, determining the molecular underpinnings of this diversity has been challenging. The transcriptional heterogeneity observed in single-cell measurements of mature neuronal cell types^5,6^ also takes place at the level of the neuronal progenitor cells (NPCs). This is evidenced by previous bulk measurements of purified progenitor subpopulations^19^ or single-cell studies in both human and mouse^20,21^. The extent of transcriptional diversity in mouse NPCs is likely to be influenced by a variety of unexamined factors that include programmed cell death^22^, genomic mosaicism^23–25^ as well as a variety of “environmental” influences such as changes in exposure to signaling lipids affecting NPCs that have been associated with disease^26–28^. We therefore used scRNA-seq to assess a proof-of-concept cohort of NPCs from an embryonic age (day 13.5) of cerebral cortical development that might exhibit transcriptional heterogeneity aspects associated with one or more of these processes. We demonstrate that PAGODA effectively recovers the known neuroanatomical and functional organization of neuronal progenitors, identifying multiple aspects of transcriptional heterogeneity within the developing mouse cortex.

## Results

### Analysis of residual variance and weighted PCA identifies informative expression features

Gene expression magnitudes vary among cells due to a combination of technical noise and biological differences in the cellular state. In a heterogeneous cell population, genes distinguishing subpopulations will exhibit higher cell-to-cell variability than expected from technical noise alone. However, accurate quantification of such excess variance (*overdispersion*) poses statistical challenges, since it is necessary to accurately account for the expected levels of technical and intrinsic biological noise^29^.

Using cell-specific error models^30^ to quantify differences in measurement quality of individual cells, we incorporated corrections for batch effects, outliers, and gene lengths, to derive robust measures of gene expression overdispersion within a given cell population. Specifically, to capture amplification and drop-out (gene detection failure) characteristics of each cell, we used the negative binomial/Poisson mixture model described previously^30^. In the original fitting process, parameters of the error model were estimated based on all pairwise comparisons between all cells. To account for the existence of multiple distinct subpopulations of cells, we modified the fitting procedure to infer the error characteristics based on comparison with *k* most similar cells (*e.g. k* = 20). The models were also extended to incorporate more flexible dependencies of the negative binomial (NB) and drop-out rate parameters (see Methods). We find that the sample variance of such NB/Poisson mixture processes can be accurately modeled as a *χ*^2^ distribution using adjusted degrees of freedom and observation weights based on the drop-out probability of a given gene (Supplementary Figure 1, see Methods).

The overdispersion for each gene was evaluated relative to a genome-wide expectation^29^. Earlier studies have shown expression magnitude to be strongly correlated with technical and, likely, biological variance in single-cell measurements^15,29,30^. n order to account for this dependence on expression magnitude, we analyzed the residual variance relative to a generalized additive model incorporating the population-average expression magnitude (Figure 1, see Methods). We find that compared to previously published approach^29^, our method robustly selects biologically meaningful sets of genes on real and simulated data (Supplementary Figure 2). Robustness can be further increased with Winsorization (Supplementary Figure 2).

**Figure 2.**
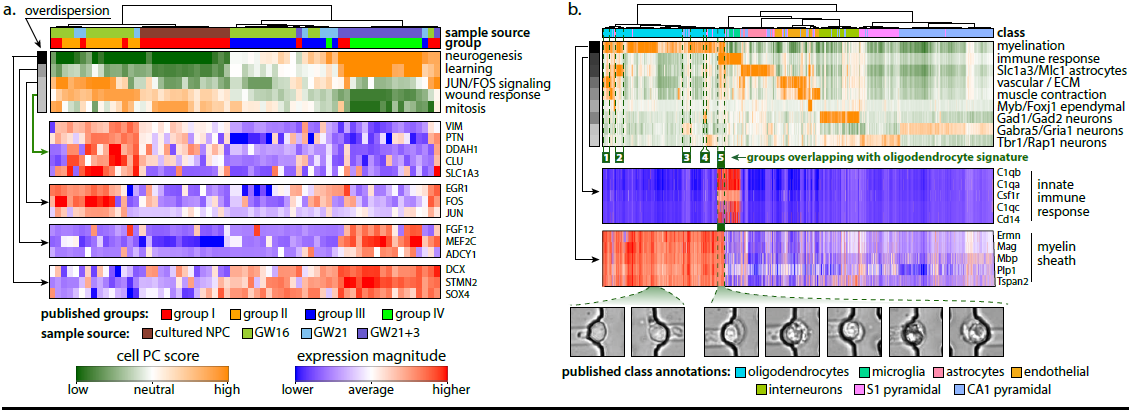
PAGODA analysis of previously published single-cell neural datasets. Transcriptional heterogeneity in the mixture of cultured human neuronal progenitor cells (NPCs) and primary cortical samples from Pollen *et al*^21^. The dendrogram shows the overall clustering of 65 cells (columns), with the top panel specifying the sample source and the group to which each cell was assigned in the original analysis by Pollen *et al*. The main panel shows five significant aspects of heterogeneity (rows) detected in the dataset by PAGODA based on gene sets defined by GO annotations, with the orange/white/green gradient indicating high/neutral/low score of a cell with respect to a given aspect. The aspect scores are oriented so that high (orange) and low (green) values generally correspond, respectively, to increased and decreased expression of the associated gene sets. Row labels summarize the key functional annotations of the gene sets in each aspect. Subsequent panels show top-loading genes underlying the identified aspects. Full composition of the identified aspects and associated expression patterns can be explored using the interactive online view^46^. Neurogenesis pathways show the strongest overdispersion within the data (Z-scores are indicated by the grayscale bar on the left), accounting for the separation of the left (progenitor) and right (maturing neurons) subpopulations. Progenitor subpopulation can be further subdivided based on activity of JUN/FOS signaling, which tracks closely with cell source (primary vs. cultured). For the “wound response” aspect, which describes a pattern complementary to the activation of neurogenesis pathway, top five genes of a *de novo* cluster driving the aspect are shown (green arrow). **b.** Top aspects of transcriptional heterogeneity in the 3005 cells from mouse cortex and hippocampus measured by Zeisel *et al*^6^. Top 9 aspects (rows) are shown, following grouping of aspects with Pearson linear correlation greater then 0.05. The aspects are labeled according to the key GO functions or the highest loading genes. Detailed composition is available through an interactive online view^46^. The detected aspects distinguish cells in a manner consistent to the top-level cell classes described by Zeisel *et al* (color code under the dendrogram). Many of the aspects overlap, indicating presence of cells exhibiting signatures characteristic of more then one cell type. Expression panels show expression patterns of top-loading genes in two GO categories: innate immune response (from the aspect distinguishing neuroglia), and myelin shealth (distinguishing oligodendrocytes). A population of ∼ 35 cells expressing both signatures is marked by a green bar, and most likely represents capture of two associated cells of different type. The bottom panel shows images of the microfluidic traps corresponding to some of the dual-signature cells, along with cells (leftmost two) exhibiting only the oligedendrocyte signature. Green boxes below the main panel highlight cells showing a combination of the oligodendrocyte signature with other cell types (numbered 1-5: vascular endothelial, astrocytes, CA1 neurons, Gad1/2 interneurons and neuroglia).

Experimental design and technical considerations commonly result in multiple batches of scRNA-seq measurements that need to be analyzed together. Batch effects are particularly problematic for heterogeneity analysis, as batch differences can produce false positive signals and obscure the presence of biologically meaningful subpopulations. To adjust for batch effects, we compared the variance estimates based on population-wide and batch-specific estimates, and down-weighted the variance of genes that exhibited batch-specific effects (see Methods). Such correction attenuates false-positive effects of batch bias (Supplementary Figure 3). It should be noted that batch effects congruent with the subpopulations of interest, such as uneven distribution of cell subpopulations between the batches, reduce statistical power and can altogether preclude statistical detection of subpopulations.

**Figure 3.**
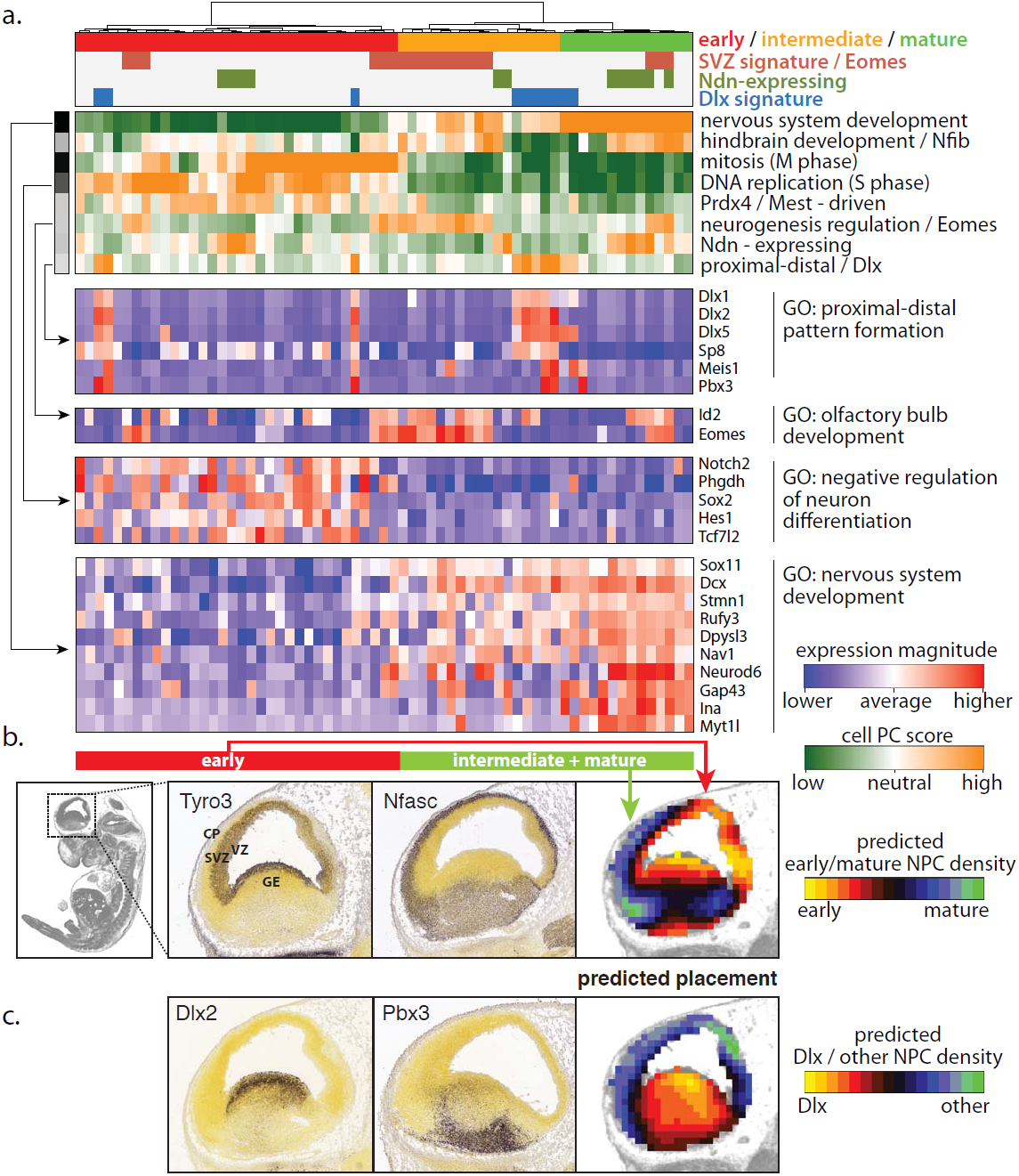
Transcriptional heterogeneity of neuronal progenitor cells in embryonic mouse cortex. **a.** Statistically significant aspects of transcriptional heterogeneity amongst E13.5 mouse cortical NPCs. Eight significant aspects that were detected following grouping of redundant patterns (Pearson r **>** 0.1) are labeled by their primary GO category or driving genes. Detailed composition is available through an interactive online view^46^. The most significant aspect (top aspect) tracks induction of neuronal maturation pathways, driving the overall subpopulation structure (separating early, intermediate and mature NPCs). Mitotic and S-phase signatures in early NPCs account for the next two most significant aspects, with the S-phase aspect incorporating closely matching expression patterns of genes responsible for NPC maintenance (see “negative regulation of neuron differentiation” expression panel below). Color codes in the top panel summarize key subpopulations of NPCs distinguished by the detected heterogeneity aspects. **b.** Anatomical placement of the early *vs.* maturing NPC classes within embryonic brain. In situ hybridization signals in E13.5 mouse brain are shown for two previously uncharacterized genes differentially expressed between early and intermediate+mature NPCs. Tyro3, upregulated in the early NPCs, is enriched near VZ (ventricular zone). Nfasc, upregulated in maturing NPCs is enriched in the CP (cortical plate) region. Computational prediction of the spatial distribution of early *vs.* maturing NPCs based on the overall transcriptional profile (third panel) places early NPCs near VZ, and maturing ones in SVZ (subventricular zone)/CP regions, consistent with known placement of apical (early) and basal (intermediate) progenitors. *In situ* images and spatial data on gene expression in the E13.5 mouse brain were generated by Allen Institute for Brain Science as part of the Developing Mouse Brain Atlas^36^. **c.** Anatomical placement of the Dlx-expressing NPCs. In situ signals for two genes upregulated in the cells exhibiting Dlx-driven signature are shown in the E13.5 mouse brain. A computational prediction places such cells in the GE (ganglionic eminence region), consistent with the anatomical origination of the tangentially-migrating NPCs.

Principal component analysis (PCA) has been widely utilized for separating subpopulations in scRNA-seq data^7,10,31^. The sensitivity of PCA, however, is limited, due to the non-Gaussian distribution of the read counts or gene expression magnitudes. While variance-stabilizing transformations – typically based on square root or log functions – can be used to improve PCA performance in such cases, the presence of drop-out events and cells with different levels of amplification noise can mask the underlying covariance structure. For instance, a subset of cells exhibiting higher drop out rates is likely to be recognized as a distinct subpopulation regardless of their biological state (Supplementary Figure 4). To account for the expected technical noise within the data, we used a weighted PCA approach^32^, weighting the individual observations based on the probability of failing to detect a transcript in a given cell due to drop-out or negative binomial dispersion. We find that this weighted PCA method exhibits significantly better performance in recovering the underlying population structure and separating individual cell subpopulations compared to the standard PCA and ICA approaches (Supplementary Figures 4, 5).

**Figure 4.**
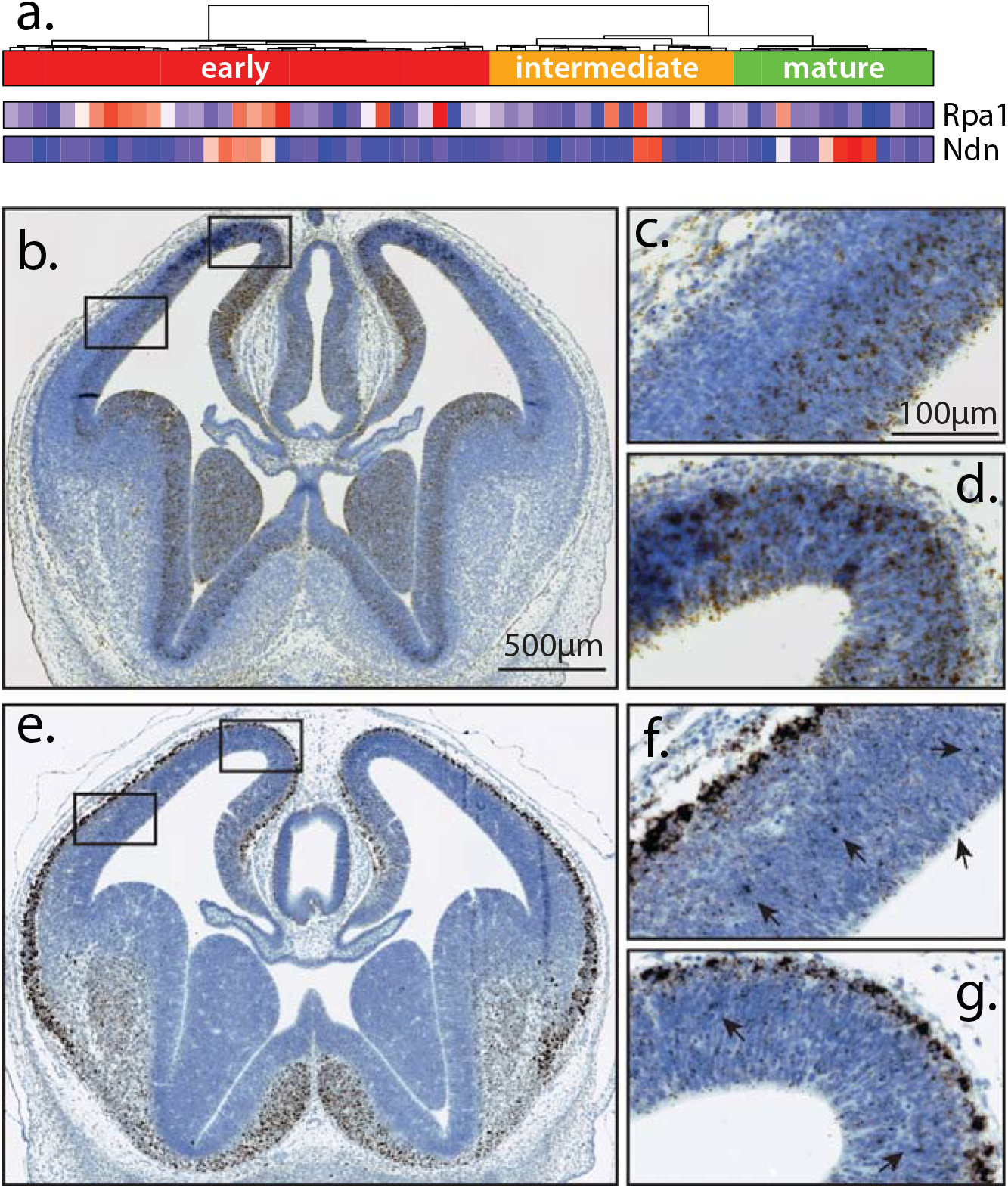
Validation of additional genes associated with NPC sub-populations identified by PAGODA. PAGODA clustering identified previously unassessed genes that would be predicted to identify distinct populations of NPCs. Two such genes, Rpa1 and Ndn, were examined by *in situ* hybridization. **a**: scRNA-seq expression magnitudes of the two selected genes relative to Figure 3a cell clustering and classification. **b-g**: Coronal E13.5 brain sections labeled by *in situ* hybridization using RNAscope probes for Rpa1, found in proliferating clusters (**b-d**) and Ndn in more mature clusters (**e-g**), representing genes with previously unknown relationship to NPCs; all sections were counterstained with hematoxylin. **b-d**: Rpa1 shows high expression in the ventricular (VZ) and sub-ventricular zone (SVZ) with reduced expression in the region of young postmitoic neurons located in the superficial cortical plate (CP). **e-g**: Ndn expression is promient throughout the CP. There are also rarer high expressing cells in the VZ and SVZ (black arrows) consistent with scRNA-seq data (**a**). These expression patterns are consistent with PAGODA identification of NPC sub-populations and demonstrate how RNA-seq can identify unique populations undefined by conventional methods.

### Pathway and gene set overdispersion analysis detects multiple aspects of heterogeneity

In order to increase statistical power to detect significant aspects of heterogeneity across the cell population, we developed an alternative approach, called PAGODA, that directly tests whether the genes in pre-defined gene sets, such as annotated biological pathways, show a coordinated pattern of overdispersion. Specifically, for each pre-defined gene set, we tested whether the amount of variance explained by the first principal component significantly exceeded genome-wide background expectation, as modeled by the Tracey-Widom F_1_ distribution^33^ with adjusted degrees of freedom (see Methods). For instance, testing gene-sets corresponding to the mouse Gene Ontology (GO) annotations on the NPC/ASC data, we identify significant overdispersion in gene sets associated with axons, synapse and nervous system development (Supplementary Table 1). Simultaneously, we also detect significant overdispersion in a number of gene sets reflective of astrocyte functions such as ion and amino acid transport, cholesterol metabolism pathways. Other overdispersed gene sets, including mitosis, DNA replication and cell division pathways, reflect cell cycle heterogeneity. Some of these gene sets represent independent aspects of heterogeneity within the population, while others such as multiple pathways related to cell cycle represent redundant aspects separating the same subsets of cells, often relying on the same key group of overdispersed genes. To provide a non-redundant view of heterogeneity structure within the dataset, the statistically significant principal components from different gene sets are clustered, and those with high similarity of gene loading or cell separation patterns are combined to form a single “aspect” of heterogeneity (Figure 1).

PAGODA allows for simultaneous detection and interpretation of the transcriptional heterogeneity within the cell population. To demonstrate PAGODA, we first examine several published datasets of varying size and population complexity. Pollen *et al.* have recently sequenced transcriptomes of 65 human neuronal lineage cells, combining cultured NPCs and primary cortical samples^21^. Testing GO annotations, we find 1274 significantly overdispersed gene sets (FDR≤ 0.05), with neurogenesis and neuron projection development being most significant (Supplementary Table 2). Clustering of redundant patterns (Pearson r > 0.5) reduces overdispersed gene sets into five main aspects of heterogeneity (Figure 2a). The neurogenesis-defined aspect dominates, separating progenitor cells (left) from maturing neurons (right), based on upregulation of DCX, SOX4, STMN2 and other known markers of neuronal maturation. This main aspect of heterogeneity is complemented by increased expression on the progenitor side of known progenitor markers such as genes from extracellular signaling and filament pathways (*e.g.* NOTCH2, SPARC, VIM). The second major aspect separates a subpopulation of more mature neurons based on sets of genes associated with learning, behavior and synapse organization. Two remaining aspects cut across this overall organization. One is driven by up-regulation of genes involved in JUN/FOS signaling (JUN, FOS, EGR1) and distinguishes a subset of progenitor cells, most of which originate from the primary cells collected at the gestational week (GW) 16, as well as a set of approximately eight maturing neurons. The other cross-cutting aspect is driven by a mitosis signature, which is most pronounced in the cultured NPCs, but is also apparent in a subset of primary progenitor cells and some of the maturing neurons, most of which originate from GW16. The overall cell clustering shows grouping consistent with the classification derived by Pollen *et al.* (groups I-IV), though assignment of some cells differs. This is particularly apparent for the four group I (early progenitor) cells that combine strong neurogenesis pathway expression with JUN/FOS signaling. The advantage of PAGODA is illustrated by its ability to detect the JUN/FOS signature as an independent aspect of heterogeneity that cuts across group I-IV partitioning.

While testing for overdispersion of pre-annotated gene sets provides increased statistical power, some aspects of heterogeneity may be driven by groups of genes that are not captured by these gene sets. To detect such aspects PAGODA also incorporates “*de novo* gene clusters” – groups of genes whose expression profiles are well-correlated within a given dataset. As the amount of variance explained by the first principal component for such gene sets will be in general higher than expected by chance, the statistical significance of overdispersion for such *de novo* clusters is evaluated using a different background model based on an extreme value distribution (see Methods). An example of genes grouped in such a *de novo* cluster is shown for the vimentin-associated (VIM) pattern in Figure 2a.

To illustrate PAGODA on a more complex population, we examined scRNA-seq data for 3005 cells from the mouse cortex and hippocampus from a recent publication by Zeisel *et al*^6^. This extensive dataset covers a variety of cell types, some of which exhibit very distinct expression signatures. Applying PAGODA and stringently reducing redundant patterns (Pearson r > 0.05, see Methods) reveals nine major aspects of heterogeneity that distinguish seven top-level classes identified by Zeisel *et al.* plus two lower-level subpopulations (Figure 2b). The most significant aspect distinguishes oligodendrocytes, which are the most numerous cells in the dataset, and are easily distinguished by strong overdispersion of myelination-related pathways. Similarly, overdispersion of immune, vascular and muscle-associated GO-annotated gene sets reveal identity of microglia, vascular endothelial, and mural subpopulations respectively. Other cell types, such as ependymal cells, or different types of neurons (interneurons, S1 and CA1 pyramidal) are distinguished by *de novo* gene set signatures, with most overdispersed genes revealing their identity (*e.g.* Gad1, Tbr1, Gabra5). We noted that aspects distinguishing many of the identified cell classes appear to overlap, most frequently with the myelination signature. For instance, a subset of ∼ 35 cells exhibits prominent expression of both immune response genes characteristic of microglia as well as genes responsible for the construction of myelin sheath (Figure 2b). Similarly, myelin-associated expression signature is observed for a subset of vascular cells, astrocytes, pyramidal neurons and interneurons (green groups 1-5, Figure 2b). These hybrid signatures most likely reflect cases in which an oligodendrocyte was captured in physical association with a second cell of a different type (Figure 2b). Such ambiguous cases account for a large fraction of discrepancies in the class assignment, but are apparent from the detected aspects.

PAGODA detects multiple, potentially independent aspects of transcriptional heterogeneity, thereby providing the means to focus on the biologically relevant features of heterogeneity, while omitting or actively controlling for undesired aspects such as technical variation or unrelated biological response. To illustrate this we analyzed scRNA-seq data for 81 differentiating mouse CD4+ T cells recently published by Buettner *et al*^12^. As detailed by Buettner *et al*, differences in the cell cycle phase account for a large fraction of the overall transcriptional variance in this dataset, masking a subtler pattern of Th2 cell differentiation in response to Il4 stimulation that involves up-regulation of Stat3 signaling, Il4ra, Il24 and other genes. PAGODA identifies the mitosis-associated signature as the most significant aspect of heterogeneity, recovering Il4ra/Il24 response and a closely aligned glycolysis aspects as third and fourth most significant patterns (Supplementary Figure 6). PAGODA also identifies heterogeneity among the examined T cells with respect to translational activity, expression of Ccl3/Ccl4 cytokines. Strikingly, the second most significant aspect of heterogeneity after cell cycle distinguishes a subset of cells expressing dozens of genes associated with dendrites, synaptic junctions, cell adhesion and organismal development (Supplementary Figure 6c). While these cells express some genes associated with T-cell endothelial migration (Cxcl12, Slit2)^34^, expression of other distinguishing genes, such as Treml4, Ceacam2 or Nav1 is atypical of T cells. Contrasting with a panel of immunological cell types we find that the transcriptome of the identified subpopulation is similar to dendritic cells, while similarity to the major Th classes (Th1, Th2, Th17, Tfh) is low despite good agreement between these bulk Th measurements and the other cells in the dataset (Supplementary Figure 7).

PAGODA provides an option to explicitly control for an extraneous aspect of heterogeneity or other non-categorical latent variables, by implementing the standard approach of subtracting the projection of the data onto a given aspect. We used such a procedure in the calculations presented above to control for the sequencing depth variation among the measured cells. In contrast, control for categorical co-variates is implemented through the batch correction procedure. To illustrate this we re-examined recent data on 622 mouse neurons from dorsal root ganglion presented by Usoskin *et al*^5^. The neurons were picked in three independent sessions that varied in their conditions, resulting in some systematic transcriptome differences. PAGODA successfully recovers the underlying neuronal taxonomy both with and without batch correction: however, omission of the batch correction step results in close clustering of neurons from the same batch, and detection of batch-specific expression patterns as statistically significant aspects of heterogeneity (Supplementary Figure 8).

### PAGODA characterizes multiple aspects of heterogeneity in mouse neuronal progenitors

Heterogeneity among NPCs is of particular interest because of its potential to influence downstream neural diversity. We used PAGODA to examine major aspects of transcriptional heterogeneity among the 65 NPCs isolated from the palium regions of an 13.5 day old embryonic mouse brain (E13.5), using dissociated whole cells that, at this age, can be readily isolated.

The most significant aspect of heterogeneity identified within the isolated NPCs reflects gradual induction of the genes associated with neuronal maturation and growth (Figure 3a, top aspect). Approximately half of the cells (right half) express Dcx, Sox11 and other known markers of neuronal maturation, with the most mature subset expressing genes involved in neuronal maturation and growth cones (*i.e.* Neurod6, Gap43). While these cells show clear commitment to the neuronal fate, they maintain expression of some progenitor markers (*e.g.* vimentin) and therefore likely represent committed NPCs.

The set of early NPCs, lacking expression of neuronal maturation markers (left half, Figure 3a), show strong up-regulation of cell cycle pathways, with M-phase and S-phase separating into two distinct aspects. These actively dividing NPCs also show up-regulation of genes characteristic of early progenitor state^20^ (Sox2, Notch2, Hes1), identified through statistically significant overdispersion of gene sets defined by “negative regulation of neuronal differentiation” and “neural tube development” GO categories. Despite the ability of PAGODA to characterize multiple aspects for a given cell, we find that neuronal maturation and S/M-phase signatures are largely mutually exclusive, with only one mitotic cell expressing genes associated with neuronal maturation (Neurod6, Dcx, Gap43, *etc*).

Maturation of neuronal progenitors is closely tied to the spatial organization of the developing cortex^35^. The early (apical) progenitors reside primarily in the ventricular zone (VZ), with intermediate (basal) progenitors occupying subventricular zone (SVZ), and maturing neurons found primarily in cortical plate (CP). Taking advantage of the extensive standardized data on the spatial expression patterns of genes in the developing mouse brain^36^, we used the expression pattern of genes differentially expressed between the early and maturing NPCs to reconstruct the most likely spatial distribution of these cells in the E13.5 mouse brain (Figure 3b, see Methods). In agreement with key marker genes, we found maturing cells to be localized in the CP and SVZ while early NPCs localize close to VZ. This spatial separation is further confirmed by examination of individual differentially expressed genes that were not previously annotated in the relevant neurodevelopmental pathways (Figure 3b).

To assess the ability of PAGODA to identify new gene relationships, we selected two genes of unknown relationship to the embryonic cerebral cortex for *in situ* hybridization analyses: Rpa1 and Ndn (necdin). Initial qPCR from bulk E13.5 brain mRNA confirmed expression of these genes, and they were therefore assessed in tissues sections using RNAscope probes (see Methods). Both genes showed robust expression with the embryonic brain, while also showing spatially distinct expression patterns that were consistent with their expression in clustered populations (Figure 4). Rpa1 was most prominent in proliferative regions. Ndn localized in the postmitotic regions (especially the cortical plate, or future grey matter of the cerebral cortex), as well as rare cells within the SVZ.

An additional subset of NPCs was distinguished by expression of Eomes (also known as Tbr2), Neurod1 and other genes whose expression has been previously shown to localize to the SVZ region and thought to distinguish basal progenitors^20,37^. Most of the cells marked by the Eomes signature express intermediate levels of genes associated with neuronal maturation (“intermediate” maturity cluster – orange, Figure 3a), however it also distinguishes a small subset of mature NPCs, as well as a subset of early NPCs undergoing DNA replication (S-phase). These likely represent neuronally-committed NPCs maturing in SVZ, and dividing basal progenitors, respectively. These dividing cells express notch signaling genes (Dll1, Notch2, Mfng) concurrently with Eomes and therefore likely represent nascent basal progenitors^20^.

Two other aspects cut across the main NPC maturation axis. The first is driven by prominent expression of necdin (Ndn) in a subset of early, intermediate and mature NPCs (Figure 3a). Necdin, initially noted for high expression in mature neurons^38^, has also been shown to be expressed in the VZ^39^. Necdin has been shown to restrict both proliferation and apoptosis rates in NPCs^39,40^, by repressing activation of E2F1 targets, including Cdk1^41^. In combination with RNAscope microscopy (Figure 4d-g), our results show that in addition to mature neurons, necdin is expressed within a subset of NPCs, approximately a quarter of which exhibit pronounced mitotic signatures and are likely localized in the SVZ.

The second notable cross-cutting aspect is characterized by coordinated expression of several Dlx homeodomain transcription factors, identified through overdispersion of gene sets defined by the “proximal/distal pattern formation” and “sequence-specific transcriptional regulation” GO categories. Dlx1/Dlx2 label tangentially-migrating NPCs, which originate in the ganglionic eminence (GE) and migrate to the cortical areas, giving rise to the GABAergic neurons^42,43^. The Dlx-positive cells express other markers of tangentially migrating NPCs, most notably Sp9 and Sp8 transcription factors^44^. Indeed, spatial localization of these cells based on their overall transcriptional profile was predicted to be in the GE region, where tangentially-migrating NPCs are thought to originate (Figure 3c). In agreement with earlier observations of such NPCs undergoing mitosis in the cortical VZ/SVZ areas, we find that out of ten Dlx-positive NPCs, two were captured in S-phase and one in M-phase. Although coordinated expression of Dlx and necdin has been implicated in differentiation of GABAergic neurons^45^, we found no overlap between the Dlx and necdin-driven aspects in the examined NPCs.

## Discussion

Just like organisms as a whole, individual cells can be classified according to a variety of meaningful criteria. Our results show that the detailed snapshots of the transcriptional state provided by the scRNA-seq often allow one to decouple multiple biological processes distinguishing a given cell within the measured population. This is seen in the case of the tangentially migrating NPCs that, despite being a distinct progenitor subtype, go through the same neuronal maturation process as other cortical NPCs, or in the case of the two distinct Th2 subsets that are also going through the common cell cycle stages. By identifying significantly overdispersed gene sets, PAGODA is able to effectively recover such complex heterogeneity structures in a wide range of datasets. In addition to increasing statistical power, the analysis of curated gene sets, such as Gene Ontology categories utilized here, also provides important clues about the likely functional interpretation of the detected aspects of transcriptional heterogeneity. To further facilitate interpretation of the heterogeneity structure PAGODA implements an interactive web browser interface^46^ for examining gene sets underlying the detected aspects of heterogeneity, and relevant expression patterns.

The potential ambiguity of the population structure also suggests an iterative approach: if, after an initial round of analysis, some of the identified aspects can be attributed to technical or biological variation irrelevant for the biological context being investigated, the expression data can be reanalyzed after explicitly controlling for such variation. Similarly, if the initial assessment of heterogeneity structure in the entire measured population identifies a prominent subpopulation of cells, PAGODA can be applied to this subset of cells, which will increase statistical power to detect heterogeneity within that subpopulation.

Our examination of transcriptional heterogeneity of mouse cortical NPCs clearly recovered the expected NPC maturation progression, marked by gradual induction of genes involved in neuronal maturation (Sox11, Dcx, Neurod6), accompanied by loss of signatures associated with maintenance of NPCs (Sox2, Hes1) and cell cycle progression^20^. Our results also illustrate a variety of additional characteristics distinguishing subsets of NPCs independent of their maturation stage, including Dlx2, Ndn and Eomes-associated signatures. We note, however, that in both our set of 65 NPCs and in prior datasets, some genes known to be expressed transiently – such as lysophospholipid receptor genes^47,48^ – and genes associated with cell death such as caspase-3^49,50^ – were not well represented despite their known expression and proven functions. This deficiency likely reflects technical aspects of current scRNA-seq measurements, and underscores the need for further characterization of transcriptomic diversity amongst NPCs and other neuronal populations. Application of computational techniques such as PAGODA and improved scRNA-seq protocols will help to uncover and interpret the multitude of specialized cellular subtypes and states comprising the mammalian cortex and other complex tissues.

## Methods

### Isolation and single-cell RNA-seq of mouse neural progenitor cells (NPC) and astrocytes (ASC)s

Single NPCs were isolated from embryonic day 13.5 cortices for RNA-sequencing. Timed-pregnant mice were sacrificed by deep anesthesia followed by cervical dislocation. The embryos were quickly removed and cortical hemispheres were isolated, ganglionic eminences removed, and all pups brains were pooled. All animal protocols were approved by the Institutional Animal Care and Use Committee at The Scripps Research Institute (La Jolla, CA) and conform to the National Institutes of Health guidelines.

Single cells were isolated by gentle trituration in ice-cold phosphate buffered saline containing 2 mM EGTA (PBSE) using P1000 tips with decreasing bore diameter. Cells were then filtered through a 40 uM nylon cell strainer and stained with propidium iodide (PI), a live-dead stain, and fluorescence activated single cell sorting (FACS) was performed selecting for PI negative cells. Samples remained on ice throughout the process and total processing time from cervical dislocation to sorting was limited to 2 hours. Single cells were sorted directly into cell lysis buffer provided in the Clontech SMARTer(r)^®^ Ultra(tm)™ Low RNA Kit for Illumina(r) Sequencing (cat # 634936), and sequencing libraries were generated using the manufacturer’s protocol. Resulting libraries were sequenced on the Illumina(r) HiSeqTM 2000 sequencing platform.

### Gene validation using *in situ* hybridization with RNA-scope

Mouse E13.5 embryos were removed from timed pregnant mice and prepared according to RNAscope instructions for paraffin embedded tissue. RNAscope probes (Advanced Cell Diagnostics) were designed by the manufacturer (Cat. #: GINS2 435891, RPA1 435911) and sections were processed using RNAscope 2.0 High Definition Reagent Kit -BROWN (Cat. #:310035) according to the manufacturer’s instructions. Sections were imaged on a Ziess Axioimager at 20X magnification.

### Previously published single-cell RNA-seq data

For the mixture of cultured human neuronal progenitor cells (NPCs) and primary cortical samples from Pollen *et al*^21^, SRA files for each study were downloaded from the Sequence Read Archive (http://www.ncbi.nlm.nih.gov/sra) and converted to FASTQ format using the SRA toolkit (v2.3.5). FASTQ files were aligned to the human reference genome (hg19) using Tophat (v2.0.10) with Bowtie2 (v2.1.0) and Samtools (v0.1.19). Gene expression counts were quantified using HTSeq (v0.5.4). Read counts for the Th2 data by Buettner *et al*^12^ were downloaded from the supplementary site (http://github.com/PMBio/scLVM/blob/master/data/Tcell/data_Tcells.Rdata). Read (or UMI) count matrices for other two datasets were downloaded from GEO: GSE60361 for Zeisel *et al*^6^; GSE59739 for Usoskin *et al*^5^.

### Fitting single-cell error models

Following the approach described in Kharchenko *et al*^30^, the read count for a gene *g* in a cell *i* was modeled as a mixture of a negative binomial (signal) and Poisson (drop-out) components: 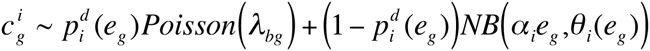, where 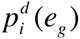 is the probability of encountering a drop-out event in a cell *i* for a gene with population-wide expected expression magnitude *eg* (FPKM); λ*bg* = 0.1 is the low-level signal rate for the dropped-out observations; θ*i* (*eg*) is the negative binomial size parameter (see functional form below); and α*i* is the library size of cell *i*, as inferred by the fitting procedure. The single-cell error models were fitted using the approach described in Kharchenko *et al*^30^, with the following modifications. **1.** Rather than estimating expected expression magnitudes of genes using all pairwise comparisons between all other cells, each cell was compared to its *k* most similar cells (based on Pearson linear correlation of genes detected in both cells for any pair of cells). The value of *k* was chosen to approximate the complexity of the dataset (1/3^rd^ of the cells for mouse and human NPC datasets, 1/5^th^ for the larger Zeisel *et al.*^6^ and Usoskin *et al.*^5^ datasets). **2.** The count dependency on the expected expression magnitude was estimated on the linear scale with zero intercept. **3**. To improve fit, the drop-out probability was modeled using logistic regression on both expression magnitude (log scale) and its square value. **4.** Instead of fitting a constant value for the negative binomial size parameter *θ*, it was fit as a function of expression magnitude, using the following functional form: log(*θ*) = *a* + *h* /(1+10^(*x* – *m*)\**s*^)^*r*^, where *x* is the expression magnitude (log scale), and *a,h,m,s,r* are parameters of the fit. This functional form provides a more flexible fit than the *θ* = (*a*_0_ + *a*_1_ / *x*)^−1^ form used in DESeq^51^, while allowing for stable asymptotic behavior.

### Evaluating overdispersion of individual genes

For each gene, the approach estimates the ratio of observed to expected expression variance and the statistical significance of the observed deviation from the expected value. To illustrate the rationale, we start with a Poisson approximation. Let 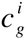 be the number of reads observed for a gene *g* in a cell *i*. If such reads follow a Poisson distribution with the mean μ*g* and variance *vg* (both equal to some Poisson rate λ*_g_*), then Fisher’s index of dispersion 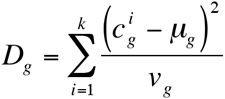 follows 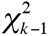 distribution^52^. While for the Poisson case *v*_*g*_ = *μ*_*g*_, for negative binomial process, *v* = *μ*_*g*_ + (*μ*_*g*_)^2^ /*θ*, where *θ* is the size parameter. As *θ* decreases from very high values where the negative binomial is well approximated by a Poisson, *D*_*g*_ diverges from 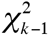. Analytical adjustments of *D*_*g*_ based on the negative binomial moments can improve *χ*^2^ approximation^53^. For more accurate approximation we used a numeric correction of the *χ*^2^ degrees of freedom, depending on the magnitude of *θ*, so that 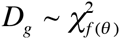 (Supp. Figure 1).

To account for the possibility of drop-out events, weighted sample variance estimates were used, so that: 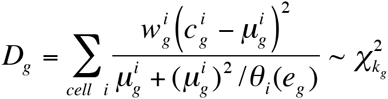, where 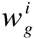 is the probability that the measurement in a cell *i* was not a dropout event based on the error model for cell *i*, and 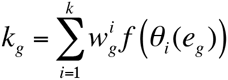 is the effective degrees of freedom for the gene *g*.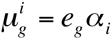, where *e*_*g*_ is the expected expression magnitude of a gene *g* across the measured cells.

Since negative binomial (or NB/Poisson mixture) models do not fully capture the variability trends observed in the real scRNA-seq measurements, *D*_*g*_ estimates for the real data can systematically deviate from 1. To adjust for this non-centrality, we normalized *D*_*g*_ by its transcriptome-wide expectation value 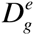, where 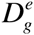 models the transcriptome-wide dependency of *D*_*g*_ on gene expression magnitude. 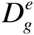 estimates were obtained using a general additive model (GAM, fit using the *mgcv* R package) as a smooth function of gene expression magnitude *eg*. To improve smoothness, the GAM fit was performed on the corresponding squared coefficient of residual variance (*D*_*g*_/*e*_*g*_)^2^. The P value of overdispersion for a gene *g* was then be calculated as 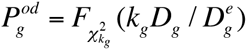, where 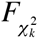 is CDF of *χ*^2^ distribution with *k* degrees of freedom.

### Weighted PCA and significance of pathway overdispersion

For PCA the data was transformed to better approximate the standard normal distribution. Specifically, PCA was carried out on a matrix of log-transformed read counts with a pseudocount of 1, normalized by the library size: 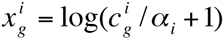. The values for each gene (matrix row) were then scaled so that the weighted variance of a given gene matched the tail probabilities of the distribution for a standard normal process: 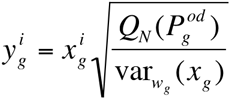, where *Q*_*N*_ is the quantile function of the standard normal distribution, and var_*wg*_ (*x*_*g*_) is the weighted variance of values *x*_*g*_. As in our previous work^30^, the weight used for the clustering and PCA steps included an additional damping coefficient *k=0.9*: 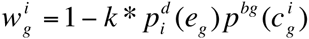, which improved the stability of the subsequent cell clustering for noisy datasets (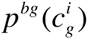 is a probability of observing 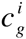 counts in a drop-out event, evaluated from the Poisson PDF).

Weighted PCA was performed for each gene set as described by S. Bailey^32^, recording first (and optionally subsequent) principal components, the magnitude of the eigenvalue (*λ*_1_) and associated cell scores for each gene set. Statistical significance of the *λ*_1_ eigenvalues obtained for each gene set (overdispersion P value for a set 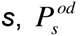) was evaluated based on the Tracy-Widom F_1_ distribution^33^ *F* (*m*,*n*_*e*_), where *m* is the number of genes in a given set *s*, and *n*_*e*_ is the effective number of cells, determined to fit the distribution of the randomly sampled gene sets (containing the same number of genes as the actual gene sets).

### Clustering of redundant heterogeneity patterns

To compile a non-redundant set of aspects, the PC cell scores (projections on the eigenvector) from each significantly overdispersed (5% FDR) gene set were normalized so that the magnitude of their variance corresponds to the tail probability of the *χ*^2^ distribution: 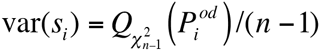, where 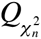 is the quantile function of the *χ*^2^ distribution with *n* degrees of freedom (*n* is the number of cells in the dataset). The redundant aspects of heterogeneity were reduced in two steps. First, aspects reflecting transcriptional variation of the same genes were grouped by evaluating similarity of the corresponding gene loading scores in combination with the pattern similarity using the following distance measure between gene sets *i* and 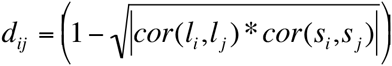 where *cor* is Peason linear correlation, *l*_*i*_,*l*_*j*_ are the loading scores of genes found in both *i* and *j* sets, and *s*_*i*_,*s*_*j*_ are the corresponding PC cell scores (*d*_*ij*_ was set to 1 if there were less than 2 genes in common between the gene sets *i* and *j)*. The distance *d*_*ij*_ was then used to cluster the aspects, using hierarchical clustering with complete-linkage. Clusters separated by a distance less then 0.1 were grouped. The cell scores of the grouped aspects were determined as cell scores of the first principal component of all aspects within a grouped cluster. The second step, aimed at grouping aspects showing similar patterns of cell separation, was accomplished by another round of hierarchical clustering using *cor*(*s*_*i*_,*s*_*j*_) distance measure with Ward clustering procedure. The similarity threshold for the final grouping of similar aspects varied between datasets depending on their complexity (0.5 for the human NPC data, 0.95 for the mouse cortical/hippocampal dataset, 0.9 for the T cell and the mouse NPC data).

### Batch correction

To control for the effect of categorical covariates, such as presence of multiple batches in the data, the approach contrasted whole-population and batch-specific variance estimates. Specifically, for each gene *g*, a batch-specific average expression magnitude was estimated for each batch *b*: *e*_*g,b*_. These batch-specific expression estimates were then used to obtain batch-adjusted values of *D*_*g*_, 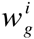 and *k*_*g*_ (*D*_*g,b*_, 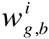 and *k*_*g,b*_ respectively). To identify genes showing batch-specific variation, the ratio of batch-specific and batch-adjusted variance was evaluated as *α*_*g*_ = *D*_*g.b*_ / *D*_*g*_. The residual variance of genes showing discrepant batch-and population-specific variance was taken to be 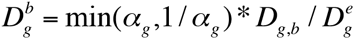, and 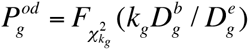.

The procedure above ensures that batch-specific effects are not reflected in the magnitude of the adjusted variance. Batch effects also need to be controlled at the level of expression values on which weighted PCA is performed, as batch-specific expression patterns across a sufficiently large set of genes can still account for sufficiently high amount of total variance to be picked by the PCA analysis. The expression values, 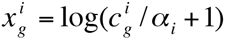, were adjusted in two steps, separating drop-out (0 read count) observations from the rest. To adjust for the disparity in the frequency of the drop-out observations between batches, the lower bound of the zero-count observation fraction (*u*) was determined for each batch (assuming binomial process), and the weights 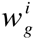 for each batch were multipled by min(1,max(*u*) / *z*_*b*_), where max(*u*) is the maximum lower bound value among batches, and *z*_*b*_ is the fraction of zero-count observations in a given batch. This procedure ensures that the expected number of zero-count observations is equal among all of the batches. The second step adjusted the log expression magnitudes of non-zero observations so that the weighted means within each are each equal to the population-wide weighted mean. To further control for batch-specific effects, weighted PCA was performed using batch-specific centering (*i.e.* setting weighted mean of each batch to 0).

### Spatial placement of cell subpopulations

To spatially place neuronal subpopulations identified by PAGODA, we used significantly differentially expressed genes (absolute corrected Z-score **>** 1.96) as relative gene expression signatures for each subpopulation of interest compared to all other NPCs. In situ hybridization (ISH) data for the developing 13.5 day embryonic mouse were downloaded from the Allen Developing Mouse Brain Atlas (Website: (c)2013 Allen Institute for Brain Science. Allen Developing Mouse Brain Atlas: http://developingmouse.brain-map.org) for all available genes (n=2194). ISH data are quantified as gene expression *energies*, defined as expression intensity times expression density, at a grid voxel level. Each voxel corresponds to a 100 µm gridding of the original ISH stain images and corresponds to voxel level structure annotations according to the accompanying developmental reference atlas ontology. The 3-D reference model for the developing 13.5 day embryonic mouse derived from Feulgen-HP yellow DNA staining was also downloaded from the Allen Developing Mouse Brain Atlas for use as a higher resolution reference image. Energies for genes in each subpopulation’s gene expression signature with corresponding ISH data available were weighted by expression fold change on a log_2_ scale and summed to constitute a composite overlay of gene expression. Background signal and expression detection in regions not annotated as part of the mouse embryo in the reference model were removed by applying a minimum gene energy level threshold of 8 units. We focused on spatial placements within the developing mouse forebrain and thus restricted gene energies to voxels annotated as ‘forebrain’ or ‘ventricles, forebrain’ in the reference atlas ontology.

### Implementation and data availability

The PAGODA functions are implemented in version 1.3 of *scde* R package, available at http://pklab.med.harvard.edu/scde/. The source code is available on GitHub (https://github.com/JEFworks/scde/). The spatial mapping of neural cells based on the data generated by the Allen Institute for Brain Science has been implemented as a separate R package, called *brainmapr*, available from GitHub (https://github.com/JEFworks/brainmapr). The single-cell RNA-seq data for the NPC and ASC cells is available from Gene Expression Omnibus (GEO) under the GSEXXXX accession number.

